# Q-score as a reliability measure for protein, nucleic acid, and small molecule atomic coordinate models derived from 3DEM density maps

**DOI:** 10.1101/2025.01.14.633006

**Authors:** Grigore Pintilie, Chenghua Shao, Zhe Wang, Brian P. Hudson, Justin W. Flatt, Michael F. Schmid, Kyle Morris, Stephen K. Burley, Wah Chiu

## Abstract

Atomic coordinate models are important in the interpretation of 3D maps produced with cryoEM and sub-tomogram averaging in cryoET, or more generically, 3D electron microscopy (3DEM). In addition to visual inspection of such maps and models, quantitative metrics convey the reliability of the atomic coordinates, in particular how well the model is supported by the experimentally determined 3DEM map. A recently introduced metric, Q-score, was shown to correlate well with the reported resolution of the map for well-fitted models. Here we present new statistical analyses of Q-scores based on its application to ∼10,000 maps and models archived in EMDB and PDB. Further we introduce two new metrics based on Q-score: Q-relative-all and Q-relative-resolution to compare a map and model to all entries in the EMDB and those with similar resolution respectively. We also explore through illustrative examples of proteins, nucleic acids, and small molecules how Q-scores can indicate whether the atomic coordinates are well-fitted to 3DEM maps and whether some parts of a map may be poorly resolved due to factors such as molecular flexibility, radiation damage, and/or conformational heterogeneity. Lastly, we show examples of how Q-scores can effectively be converted to atomic B-factors. These analyses provide a basis for how Q-scores can be interpreted effectively to evaluate 3DEM maps and atomic coordinate models prior to publication and archiving.

**Synopsis:** Q-scores are calculated each atom in models fitted to 3DEM (3D electron microscopy) maps. They measure how well the model fits the map, and also reflect the quality of the map as they correlate to resolution. Here we develop a statistical model for Q-scores applied to many maps and models in the EMDB (Electron Microscopy Database) and PDB (Protein Data Bank) respectively, and show how it can be used to assess the reliability of entire models as well as their subcomponents.

## 1. Introduction

Atomic coordinate models derived from 3DEM maps give many insights into the structure and function of biological macromolecules. Building models into 3DEM maps can take various paths, such as fitting of known models obtained previously with experimental methods (Pintilie & Chiu, 2012) or predicted with computational methods like AlphaFold (Jumper *et al*., 2021). Alternatively, in ∼3.5Å and better resolution maps, models can be built de-novo either interactively (Casañal *et al*., 2020) or automatically (Jamali *et al*., 2024). The quality of the 3DEM map can vary locally (Vilas *et al*., 2020), and it has become more critical to quantitatively assess the reliability of models and their various molecular components, which can be accomplished by the application of map-model metrics.

An example of a map-model metric is atom inclusion, an early metric which is still used in validation reports for depositions to the Electron Microscopy Data Bank (EMDB) (Joseph *et al*., 2017; wwPDB Consortium, 2024). Other map-model metrics include cross-correlation (Klaholz, 2019), mutual information (Vasishtan & Topf, 2011), EM-Ringer (Barad *et al*., 2015), and FSC-Q (Ramírez-Aportela *et al*., 2021). A recent Community Challenge in which many worldwide groups participated has compared such metrics, showing some similarities and correlations amongst them (Lawson *et al*., 2021). For example, Q-scores were shown to correlate with the overall map resolution, and hence relates to resolvability, although this also depends on whether a model is optimally fitted to the map (Burley *et al*., 2022; Pintilie *et al*., 2020).

While Q-scores have already been added to validation reports for maps and models deposited to EMDB (Kleywegt *et al*., 2024), we continue to evaluate how they may be interpreted in several contexts. For example, Q-scores can be averaged over all atoms in an entire model, in individual protein residues and nucleotides, or in small molecules such as ligands, saccharides and lipids. Here we show how Q-scores can be interpreted based on statistics derived from ∼10,000 map/model combinations available publicly in EMDB and PDB.

In particular, we carry out a comprehensive statistical analysis of how Q-scores are related to reported resolution, based on ∼10,000 maps and models archived in EMDB. The purpose of this study is to establish statistically sound metrics useful for evaluating 3DEM maps and models of biomolecules, including proteins, nucleic acids, and small-molecule ligands. As Q-scores are already included in wwPDB validation reports, another goal is to provide new percentile-based formulations to be used in such a context. The percentile Q-score based metrics introduced here compare a map and model to other 3DEM maps and models in the EMDB, and aim to serve as an indication of overall map and model quality (Gore *et al*., 2017; Feng *et al*., 2021).

We also further explore the use of Q-scores to derive atomic B-factors. Atomic B-factors have been commonly used in macromolecular crystallography (MX), and are also known as Debye-Waller Factors (Winn *et al*., 2001) or Atomic Displacement Parameters (Afonine *et al*., 2018). In 3DEM, the term B-factor is also used to describe the overall decay of high frequency information due to electron microscope parameters and detector performance factors (Rosenthal & Henderson, 2003), and also to report the amount of sharpening applied to a map to improve visualization in real-space (Kaur *et al*., 2021). Here we use the term “atomic B-factor” to distinguish their application to individual atoms in models fitted to 3DEM maps. In the field of 3DEM, atomic B-factors can be calculated during model refinement (Afonine *et al*., 2018; Beton *et al*., 2024) or molecular dynamics flexible fitting (Frank, 2017). We showed previously that atomic B-factors can also be derived from Q-scores (Zhang, Pintilie *et al*., 2020; Pintilie & Chiu, 2021). Here we expand this analysis with more examples, showing that atomic B-factors can be derived from Q-scores at resolutions ranging from ∼1 to ∼4 Å, closely reflecting the 3DEM map they are based on.

## 2. Q-scores of maps and models in EMDB and PDB

Q-scores were calculated for 10,189 map/model combinations in the EMDB and PDB, selecting primarily maps with reported resolution between 1 and 10 Å using the gold-standard FSC_0.143_ criterion (Henderson *et al*., 2012). The Q-score averaged over all non-hydrogen atoms in a model is plotted against the reported resolution in Figure 1A. A regression of these data points, using a 3rd degree polynomial (Figure 1) shows good correlation, with R^2^=0.7039. Residual plots in Supplementary Figure S1 confirm that this relationship fits the data well. We used a 3rd degree polynomial because it fits the data better with higher R^2^ than do linear (R^2^=0.5959) or 2nd degree polynomial (R^2^=0.6999) regressions, while not overfitting the data. Using a 4th degree polynomial did not significantly improve the fit (R^2^=0.7061). The 3rd degree polynomial model was also verified as the optimal polynomial regression calculation by cross-validation and visual inspection of regression residual plots.

**Figure 1.**
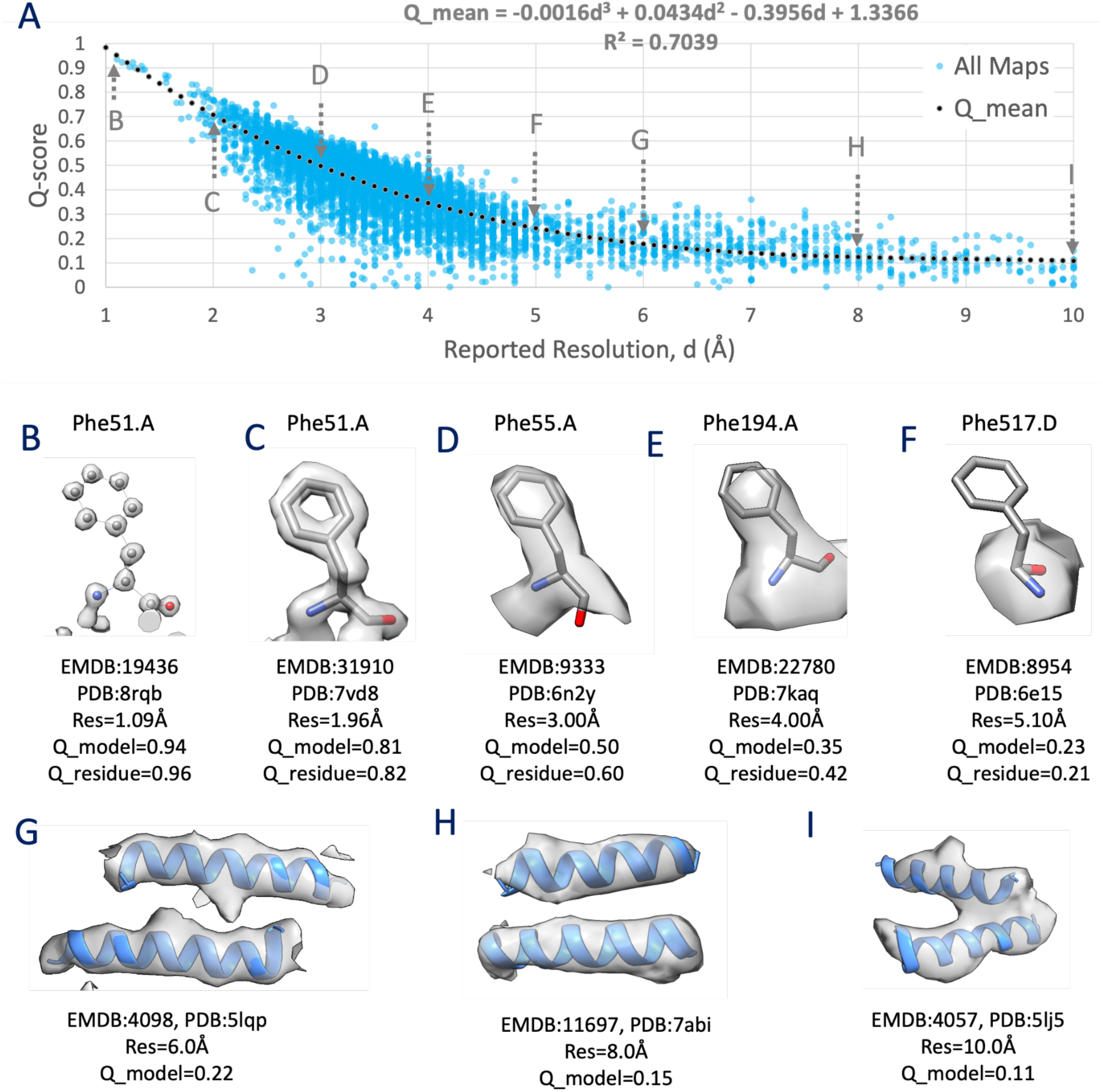
Relationship between Q-score and reported resolution, d, using EMDB maps and their associated atomic models in the PDB. (A) A plot showing each map and model pair as a filled circle, with a dotted line showing a regression using a 3rd degree polynomial. (B-F) Sidechains at various resolutions, with corresponding decreasing Q-scores, averaged over the whole model (Q_model), or averaged over the residue shown (Q_residue). (G-I) Alpha helices at 3 different resolutions between 5 and 10Å.

The plot in Figure 1 shows that Q-scores decrease quickly from ∼1 to ∼0.3 for maps with resolutions of 1 to 5 Å, and they decrease more slowly from ∼0.3 to ∼0.1 for maps with resolutions of 5 to 10 Å. Panels B to I in Figure 1 show examples of maps and models with average Q-scores near the regression line, illustrating that Q-scores correlate well with the resolvability of atoms and groups of atoms such as protein residues and alpha helices. For example, Q-scores near ∼1.,0 are associated with individually resolved atoms (Figure 1 panel B), and Q-scores near ∼0.5 are associated with resolved side chains in protein residues (Figure 1 panel D). Q-scores near ∼0.2 are associated with unresolved side chains but resolved secondary structures such as alpha helices in proteins (Figure 1 panels F-I).

Figure 1 shows some data points far away from the regression line, especially ones far below the line, with Q-scores close to 0, e.g. in the resolution range of 2.5 to 5 Å. In Supporting Information S1, we detail how removing some of these outliers using cross-correlation scores yields similar regression curves.

## 3. Statistical model for Q-scores

In Supporting Information S2, we detail how we arrive at the following equations for characterizing Q-scores at different resolutions using the polynomial regression curve illustrated in Figure 1:

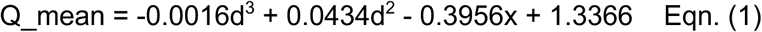

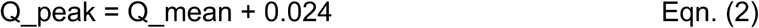

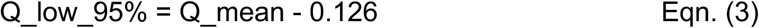

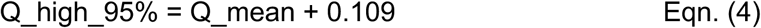

In Eqn. (1), Q_mean represents the mean Q-score value as a function of reported resolution, *d*, as calculated by regression with the 3rd degree polynomial curve illustrated in Figure 1. In Eqns. (2-4), offsets act to move the Q_mean curve up and down to three specific positions. The first is Q_peak, Eqn. (2), which positions the curve such that the highest number of data points are close to the line (within window size of 0.01). The other two positions are Q_low_95%, Eqn. (3), and Q_high_95%, Eqn. (4). These two latter offsets move the curve to positions such that 95% of the data points fall between them and Q_peak.

Q_peak represents the Q-score observed in the highest number of map-model pairs, based on the set of ∼10k maps in EMDB considered here. In statistics, this is also often called the mode of the distribution. For a normal distribution, the mean is considered to be the expected value, and coincides with the peak of the curve. In this case, because the distribution is skewed (as shown in Supplementary Figure S3B, the mean does not coincide with the peak. The other two curves, Q_low_95% and Q_high_95% provide two Q-scores below/above which a small fraction of maps (5%) are observed. Below and above these curves, Q-scores may be considered to be ‘outliers’ or ‘not commonly observed’ for a given reported resolution.

Figure 2A shows the same plot as in Figure 1A, with all ∼10k map-model pairs, also plotting the Q_peak, Q_high_95% and Q_low_95% curves. Several outliers which are outside the 95% curves are shown in Figure 2B-E. In Figure 2B and 2D, maps and models with Q-scores lower than Q_low_95% are shown. These appear to have low Q-scores due to the model not being fitted correctly to the map. Correct fitting brings the Q-scores within the 95% range.

**Figure 2.**
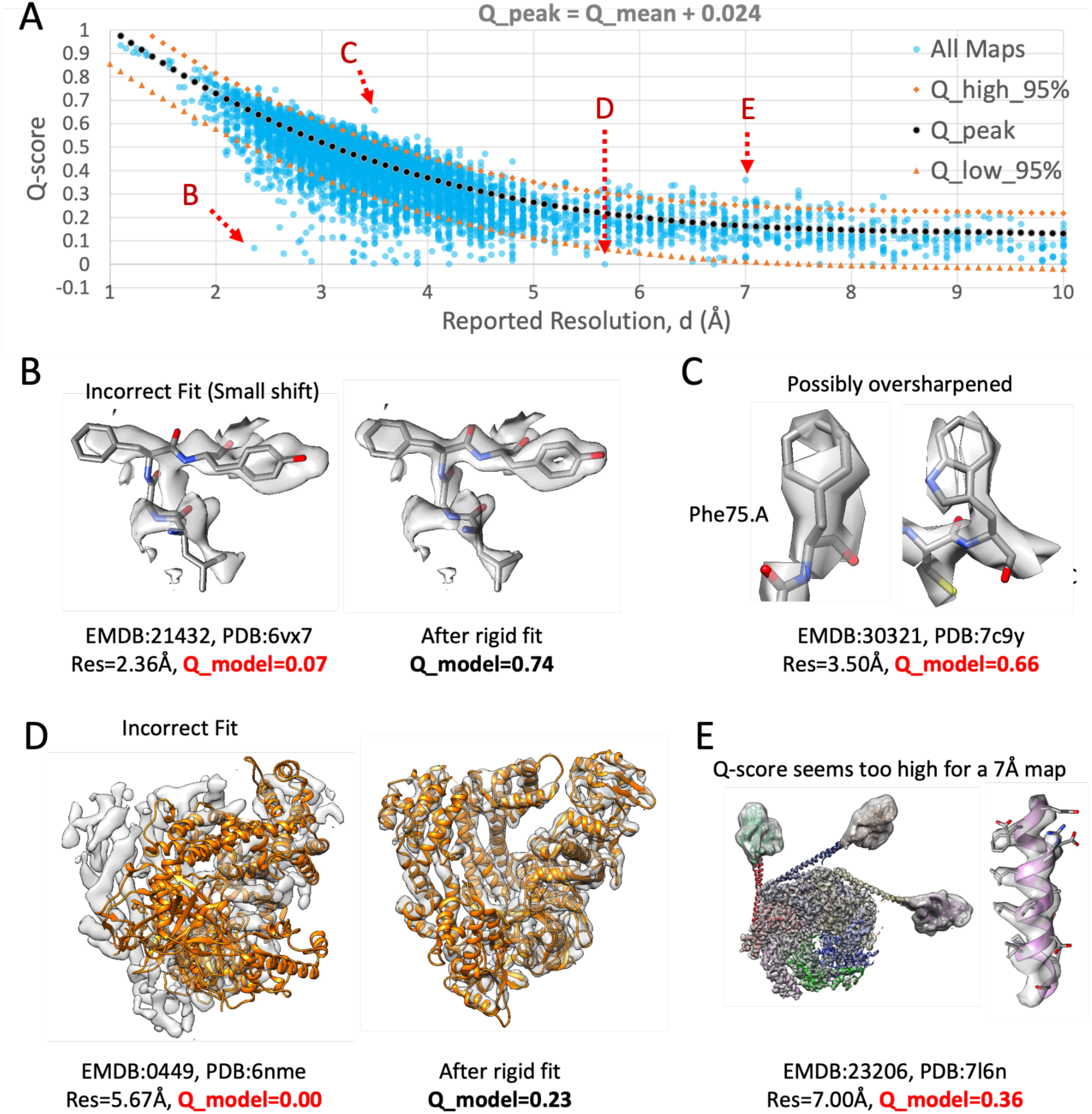
(A) Plot of Q-scores vs. reported resolution for ∼10k maps and models in the EMDB (same data set as Figure 1). The dotted curves above and below the Q_peak curve enclose 95% of the data points, Eqns. (2-4). (B-E) Illustration of maps and models with Q-scores outside the 95% curves. Overall Q-scores for each model are indicated with Q_model, color-coded red if outside the 95% curves. For B and D, Q-scores are inside the 95% curves after properly fitting the model to the map and re-calculating Q-scores.

Figure 2C shows an example where the Q-score is above the Q_high_95% line, and hence may also be considered an outlier. The map density appears discontinuous and noisy, indicating that the map is likely over-sharpened. While severe oversharpening was shown to yield lower Q-scores due to excessive noise, a small amount of over-sharpening may raise Q-scores, especially if the model is refined into the oversharpened map. Figure 2E shows another outlier where the Q-score is above the Q_high_95% curve. In this case, most of the map density seems to be resolved at higher resolution. Hence in this case the reported resolution is likely to be under-estimated and does not reflect the overall resolvability of all the features in the map.

In Supporting Information S3, we also show how we can use a rolling window approach over the same dataset, to derive similar percentile statistics without using the polynomial regression curve. The two approaches are shown to produce very similar results, however using the polynomial regression curve method appears to produce smoother curves for Q_high_95% and Q_low_95% which is advantageous.

## 4. Per-residue and Per-nucleotide Q-scores

Q-scores are calculated for each atom, but they can also be averaged over all atoms in a model (as in the previous analyses), and for groups of atoms within protein amino acid residues or nucleic acid nucleotides. We illustrate this in the examples below. Figure 3A shows a segmented map of beta-galactosidase imaged at 1.9 Å resolution, EMD-7770 (Bartesaghi *et al*., 2018). In Figure 3B, Q-scores of backbone and sidechain atoms are plotted for every residue in the associated model with PDB ID 6cvm. Q-scores of backbone atoms are mostly close to the Q_peak line calculated with Eqn. (2). Sidechain atoms however have more variable Q-scores, some below the Q_low_95 line calculated with Eqn. (3). Residues with low Q-scores for backbone and/or sidechain atoms can be identified in such a plot, as in the example shown in Figure 3B and Figure 3Aii, where low Q-scores are labeled in red. This can be used to identify areas of the map where the model may not be fitted properly, or where the 3DEM density is not well resolved and hence the accuracy of those parts of the model may be low.

**Figure 3.**
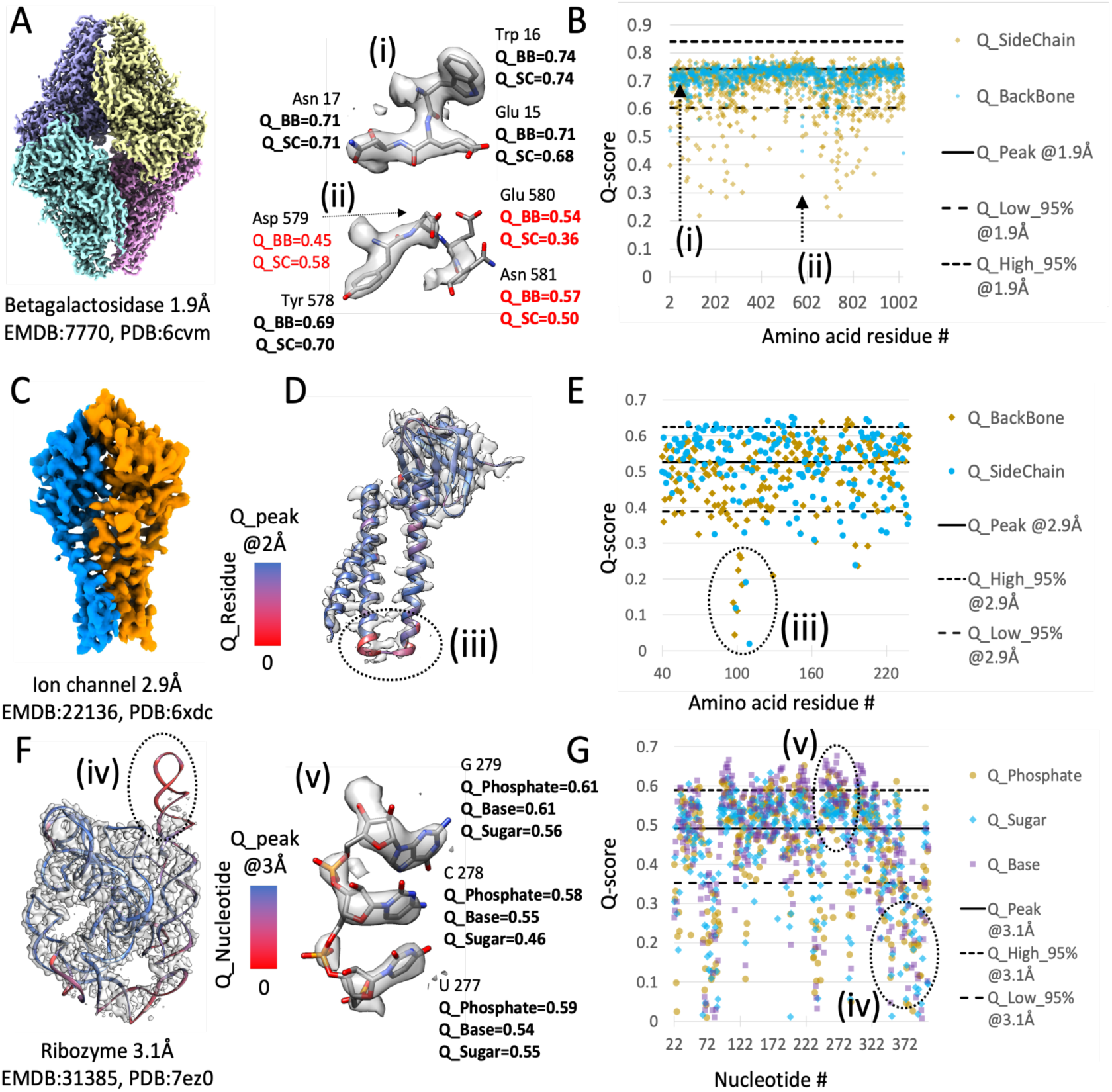
Examples of Q-score application in proteins and in nucleic acids. (A) Beta-galactosidase protein complex, with. (B) Per-residue backbone and sidechain Q-scores; example residues with Q-scores marked on the plot are marked (i) and (ii). (C) Ion channel protein complex; one of the two proteins in the complex is shown in (D), with ribbon display color coded by residue Q-score. (E) Per-residue backbone and sidechain Q-scores for one ion channel complex protein; an area with low Q-scores is marked (iii). (F) RNA-only Tetrahymena ribozyme; the ribbon model is color coded by nucleotide Q-score. (G) Q-scores of phosphate, sugar, and base atoms in each nucleotide; Q-scores for three residues which are well-resolved are shown in (v), and an area with low nucleotide Q-scores is marked (iv).

Figure 3C illustrates a 2.9 Å map of a SARS-CoV-2 ion channel, EMD-22136 (Kern *et al*., 2021). In Figure 3D per-residue Q-scores are used to color-code the backbone ribbon of one of the proteins, with red corresponding to low Q-scores (near 0), and blue corresponding to Q-scores near Q_peak (as commonly observed for this resolution). Q-scores of backbone and sidechain atoms in each residue are also plotted in Figure 3E; most fall within the 95% bounds. An area where Q-scores are much lower is marked (iii) in Figure 3E; it can also be seen as red-colored ribbon in Figure 3D, corresponding to low Q-scores. This display can be very useful for identifying areas where the map is not well resolved due to conformational heterogeneity, or where the atomic coordinate model may need further refinement to better fit the map.

Figure 3F shows a 3DEM map of the RNA-only Tetrahymena ribozyme reconstructed to 3.1 Å resolution, with id EMD:31385 (Su *et al*., 2021). Per-nucleotide Q-scores are plotted in Figure 3G. Q-scores were averaged and plotted for base, ribose, and phosphate atoms in each nucleotide. An area where Q-scores are much lower than commonly observed, under the Q_low_95 line, is marked (iv); the corresponding area in the map is not resolved well, likely due to conformational heterogeneity. An area where nucleotides are resolved as expected, and correspondingly where Q-scores are above the Q_peak line, is marked (v).

## 5. Q-scores for small molecules

Q-scores can also be calculated for small molecules, to inform whether their atomic coordinates are well resolved and/or fitted correctly in the 3DEM map. An example is a glycan, made up of smaller oligosaccharides molecules covalently bonded to proteins such as the NL63 spike trimer (Zhang, Li *et al*., 2020). In Figure 4A, a segmented 3DEM map of the coronavirus NL63 (EMD-22889) shows the three spike proteins, with ASN-associated glycans in yellow. Figure 4D plots Q-scores of each saccharide molecule. Most of the saccharide units are resolved, with Q-scores within the 95% bounds, as in the example in Figure 4B. At the same time, from the Q-score plot, it is easy to identify those that are not well resolved, as shown in Figure 4C, likely due to conformational heterogeneity.

**Figure 4.**
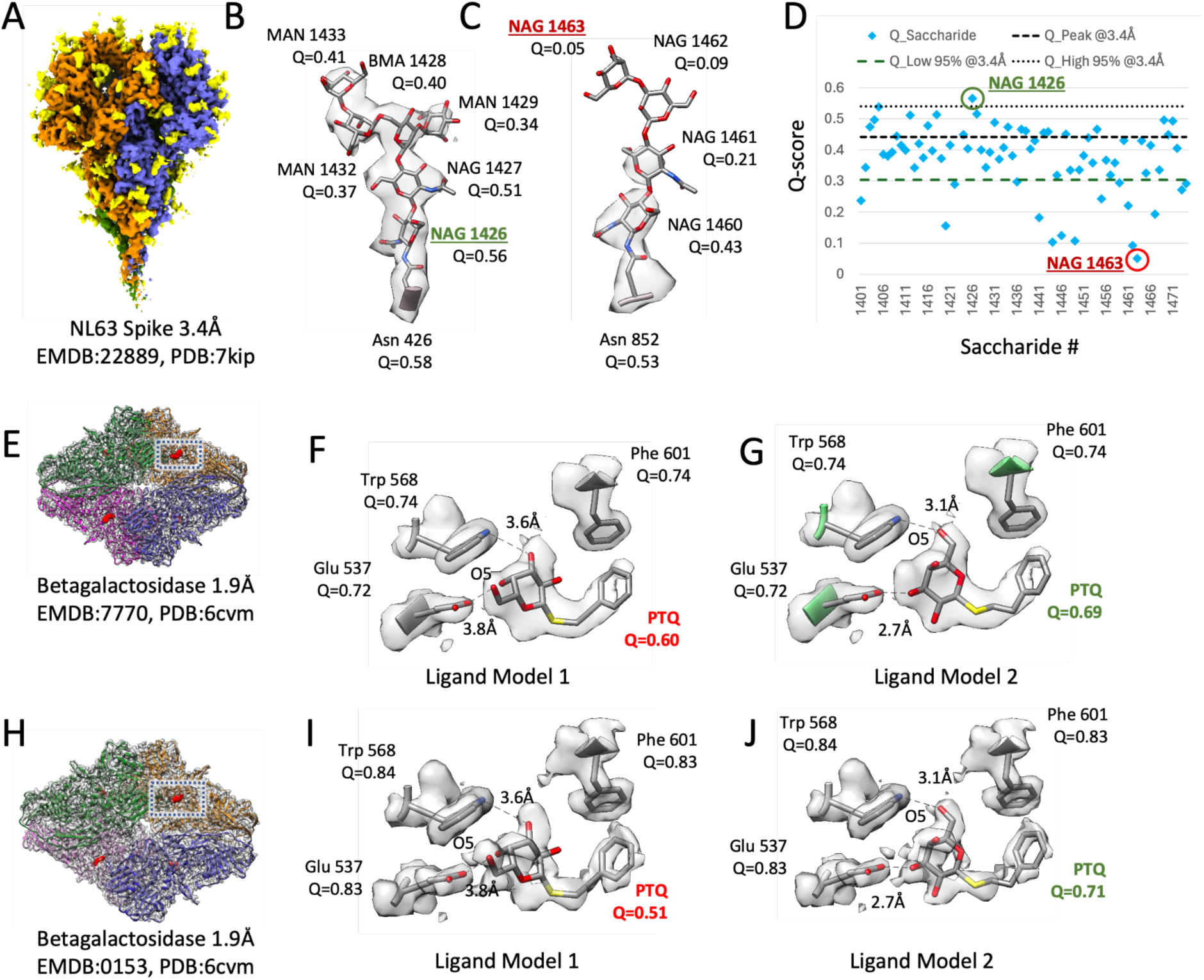
Application of Q-scores to small molecules. (A) Segmented 3DEM map of coronavirus NL63 spike proteins (blue, orange, green) with ASN-associated glycans (yellow). (B,C) Two example glycans, with Q-scores for each component saccharide. (D) Q-scores of each saccharide is plotted. (D) Q-score of each saccharide is plotted in (D). (E, H) Two 3DEM maps of beta-galactosidase with the same reported resolutions of 1.9Å. Two models of the ligand PTQ and three interacting protein residues, along with Q-scores, are shown in (F, G) for the map in (E), and in (I, J) for the map in H.

As another example, we computed Q-scores for the PTQ ligand in the beta-galactosidase complex (Bartesaghi *et al*., 2018). Figures 4E and 4H show two maps of this complex with the same reported resolution of 1.9 Å. In a recent 3DEM ligand modeling challenge (Lawson *et al*., 2024), participants reported two potential models for this ligand in the target 3DEM map EMD-7770. The two models are shown in Figures 4F and 4G. The O5 atom in the ligand is marked in both images to show the difference which is that the pyranose ring is flipped ∼180 ° in one model relative to the other. We also fitted these two models to the map of the same complex, EMD-0153, shown in Figure 4H; the fitted ligands are shown in Figures 4I and 4J. We calculated Q-scores for both ligand models in both maps. Model 1 has lower Q-scores in both maps, near or under the Q_low_95% value, and hence may be considered an outlier or unlikely. On the other hand, model 2 has higher Q-scores in both maps, in line within Q-peak or the commonly observed Q-score at this resolution; it also shows more favorable interaction distances with two nearby residues, as shown in Figures 4G and 4J. Taken together, this indicates that model 2 is more likely to be correct.

## 6. From Q-scores to B-factors

When generating a 3D map from atomic coordinates (a model-map), the effect of atomic B-factors is to spread out map values around each atom’s position. The higher the B-factor of an atom, the more diffuse or blurry, and the less sharp, the surrounding map values around the atom are. This effect can be characterized by Q-scores, because Q-scores are higher for sharper peaks, and lower for more diffuse peaks. Hence, we use a scaling parameter to calculate atomic B-factors from Q-scores, using the equation:

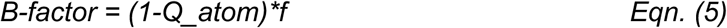

In Eqn. 5, the scaling factor *f* is determined by maximizing the similarity between the 3DEM map and the model-map generated using the resulting B-factors. Model-maps are generated with atomic B-factors resulting from scaling factors in the range of 0 to 300 and compared to the 3DEM map by cross-correlation around the mean (CC-mean). The optimal scaling factor *f* and resulting atomic B-factors are the ones that yield the highest cross-correlation score between the model-map and the 3DEM map.

Figure 5 (top row) shows residues from 4 different 3DEM maps and models with resolutions in the range of ∼1 to ∼4 Å. When using B-factors of 0 Å^2^, all residues and side chains are resolved equally (Figure 5, second row), but this does not look like the 3DEM map, where some residues are not resolved. When using B-factors derived from Q-scores, using the optimal scaling factor, the model-map looks more like the 3DEM map (Figure 5, third row) - side chains that are not resolved in the 3DEM map (and hence have low Q-scores, which would result in a high B-factors) are also not resolved in the model-map. Figure 5 (bottom row) shows plots of the CC-mean obtained with different scaling factors for each of the 4 examples, from which the optimal scaling factor (colored in orange and shown above the plot) is determined.

**Figure 5.**
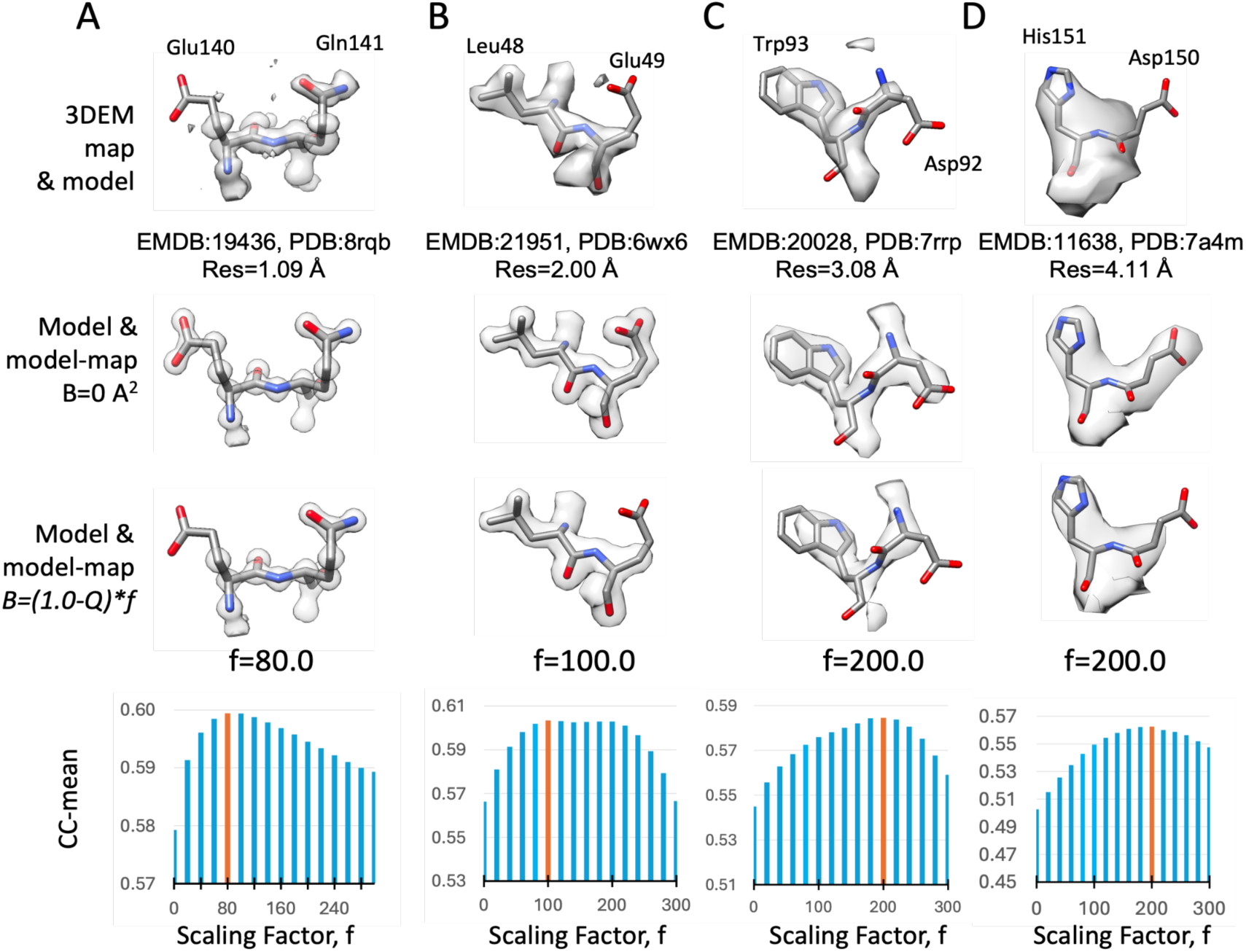
Atomic B-factors from Q-scores. Top row: two residues in 4 different 3DEM maps and models with resolutions of ∼1 to ∼4 Å. Second row: model-maps generated with atomic B-factors calculated by scaling Q-scores. Third row: model maps generated with atomic B-factors set to 0. Fourth row: bar plots of CC-mean (cross-correlation about the mean) between the 3DEM map and model-maps generated with atomic B-factors calculated using a range of scaling factors (0 - 300); the bar with the highest CC-mean value is colored orange.

## 7. Relative Q-scores

Relative Q-scores aim to compare a map-model entry to other entries in the EMDB. Here we introduce two new terms, Q-relative-all and Q-relative-resolution. Q-relative-all expresses the Q-score of a map-model entry as a percentile relative to all the entries in the EMDB, while Q-relative-resolution expresses it relative to entries with similar resolutions.

Q-relative-all is defined for a map-model pair with Q-score, Q, as follows:

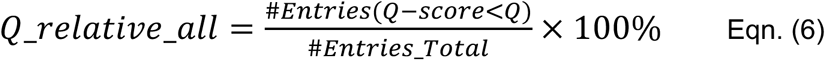

In Eqn. (6), the numerator represents the number of EMDB entries with Q-scores lower than that of the entry in question, and the denominator is the total number of entries in the EMDB. Q-relative-all thus represents the percentile ranking of an entry within the entire dataset of EMDB entries.

Q-relative-resolution is defined for a map-model pair with Q-score Q and resolution d as follows:

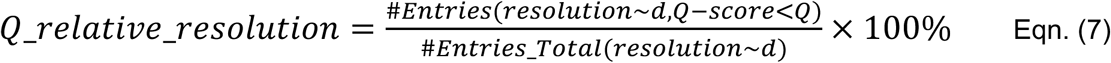

In Eqn. (7), the numerator represents the number of EMDB entries which have resolution close to *d*, more specifically within a window size *w* of the reported resolution of the entry, and also which have a lower Q-score than the Q-score of the entry, *Q*. The denominator is the total number of entries which have resolution within the same window size w of the resolution of the entry, d.

We address here what would be a good resolution window size (w) for comparing entries for calculating Q-relative-resolution. To test the effect of this resolution window size, we selected 12 window sizes ranging from 0.1 Å to 1.0 Å, with increments of 0.1 Å, including additional sizes of 1.2 Å and 1.5 Å. As shown in **Table 1**, the number of entries (minimum, mean, and maximum) increases with increasing window size. A larger number of entries for a given resolution would be more desirable, for more meaningful statistical comparison.

**Table 1.** Numbers of entries (with minimum, mean, and maximum) for different window sizes across resolutions of 1-10Å.

| Window Size, w (Å) | 0.1 | 0.2 | 0.3 | 0.4 | 0.5 | 0.6 | 0.7 | 0.8 | 0.9 | 1 | 1.2 | 1.5 |
| --- | --- | --- | --- | --- | --- | --- | --- | --- | --- | --- | --- | --- |
| MIN (# Entries) | 12 | 26 | 39 | 43 | 55 | 73 | 100 | 159 | 237 | 299 | 360 | 453 |
| MEAN (# Entries) | 2056 | 3744 | 5109 | 6761 | 8288 | 9584 | 10740 | 11791 | 12751 | 13639 | 14992 | 16364 |
| MAX (# Entries) | 3651 | 6214 | 7952 | 10282 | 12358 | 13905 | 15044 | 15922 | 16681 | 17238 | 18057 | 18698 |

A low correlation between Q-relative-resolution and reported map resolution would also be desirable, so that within each window, Q-relative-resolution is not biased towards higher reported resolution entries. Thus, we tested the correlation between Q-relative-resolution and reported resolution for different window sizes. The Pearson correlation coefficient between resolution and Q-relative-resolution is plotted in Supplementary Figure S5. Two curves are plotted, one for the correlation between Q-relative-resolution and reported map resolution, considering entries with resolution higher than 5Å (blue curve), and one considering entries with resolution lower than 5Å (red curve). For entries at resolutions lower than 5 Å, there is no significant correlation between Q-relative-resolution and reported resolution for all window sizes, as the correlation coefficient stays below 0.2 for all window sizes. However, for maps with resolutions higher than 5 Å (inclusive), negative correlations of higher magnitude are observed as the window size increases. Notably, at a window size of 0.5 Å, the correlation nears −0.3, which represents a weak degree of correlation(Evans, 1996). Thus, for little or no correlation between Q-relative-resolution and reported resolution, according to Supplementary Figure S5, the window size should be 0.5 Å or lower.

Some examples of Q-relative-all and Q-relative-resolution for maps and models presented in Figures 2 and 3 are shown in Table 2. For Q-relative-all, the higher the number, the higher the Q-score, and thus the better the overall quality of the map and model. On the other hand, Q-relative-resolution shows how the Q-score compares to other maps and models at similar resolution. The closer it is to 50%, the more it is “as commonly observed”. This would indicate a proper fit of the model to the map, and also an appropriate reported resolution value for the map. When Q-relative-resolution is much lower than this, e.g. lower than 5%, it could potentially indicate incorrect fit of the model to the map, or a map at lower resolution than reported (as shown in Figures 2B and 2D). When Q-relative-resolution it is much higher (e.g. 95% or more) it could potentially indicate other issues such as over-sharpening of the map (as shown in Figure 2C), or potentially that the reported resolution could be too low and does not reflect the overall map quality (as shown in Figure 2E).

**Table 2.**
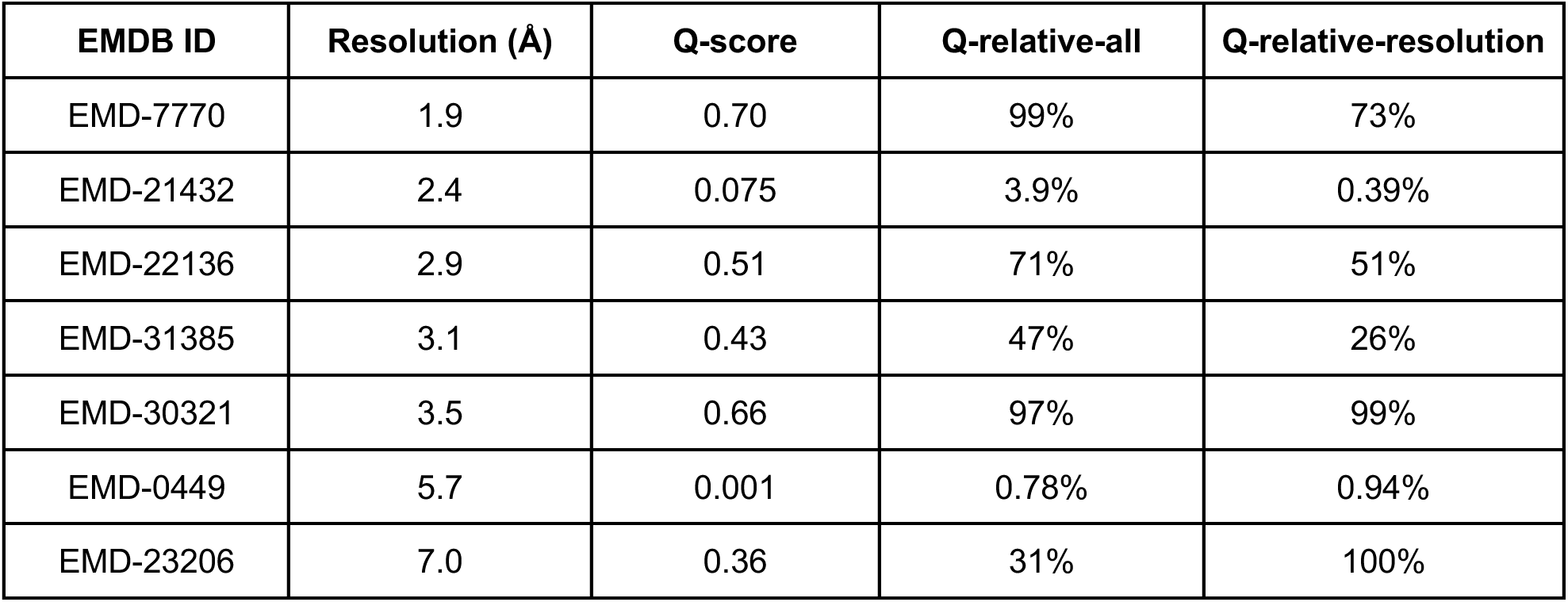
Entries from Figures 2 and 3 show corresponding Q-score, Q-relative-all, and Q-relative-resolution. The Q-relative-resolution is calculated using a resolution window size of 0.5Å.

| EMDB ID | Resolution (Å) | Q-score | Q-relative-all | Q-relative-resolution |
| --- | --- | --- | --- | --- |
| EMD-7770 | 1.9 | 0.70 | 99% | 73% |
| EMD-21432 | 2.4 | 0.075 | 3.9% | 0.39% |
| EMD-22136 | 2.9 | 0.51 | 71% | 51% |
| EMD-31385 | 3.1 | 0.43 | 47% | 26% |
| EMD-30321 | 3.5 | 0.66 | 97% | 99% |
| EMD-0449 | 5.7 | 0.001 | 0.78% | 0.94% |
| EMD-23206 | 7.0 | 0.36 | 31% | 100% |

## 8. Summary and Discussion

We previously showed that Q-scores correlate to reported resolutions of a 3DEM maps for a small but representative number of maps and models (Pintilie *et al*., 2020; Burley *et al*., 2022). Here, we further expanded the data set to ∼10k maps and models at resolutions between 1 and 10Å in the EMDB/PDB. We found that Q-scores correlate similarly to the reported resolution for this larger data set. Moreover, the distribution is close to normal, but slightly skewed towards lower Q-scores, likely due to some models not being optimally fitted to the corresponding maps. We derived a statistical model which provides, for a given resolution, the most commonly observed value, Q_peak, and also 95% bounds Q_low_95% and Q_high_95%. The latter can be used to evaluate whether a calculated Q-score is as commonly observed or instead is more of an outlier if it is outside of the 95% bounds.

We showed how this statistical model can be used to evaluate Q-scores for entire models, and for smaller groups of atoms. Such groups of atoms can be individual protein residues (also backbone and sidechain atoms), nucleic acid nucleotides (also sugar, base, or phosphate atoms), and other small molecules such as saccharides in glycans, and ligands bound to proteins. Q-scores of such groups of atoms can indicate whether protein residues, nucleotides, or small ligands are resolved as expected, and whether they are fitted optimally to the map. In the case of ligands, where more than one conformation may appear plausible, as in the example shown here, Q-scores can help to identify which conformation is more likely to be correct. Q-scores can also serve as a quantitative measure to support models of such ligands, besides relying on visual analysis alone.

In the 5 to 10 Å resolution range, we saw that Q-scores decline more slowly as a function of resolution (Figures 1,2). Thus the Q-score is less useful in this range as it is not as sensitive to the resolution of the map. However, Q-scores can still be consulted for such cases to indicate potential issues. For example, a Q-score close to 0 can suggest that the model is not properly fitted to the map, as was seen in the example in Figure 2D. We also saw an example where the Q-score for a 7 Å map was much higher than the commonly observed value (Figure 2E). Visual inspection revealed that the map contained areas of higher resolution, so the reported resolution was not fully representative of the entire map. Thus, the current formulation of Q-scores may be, for the time being, also useful in this resolution range as a means of identifying such inconsistencies.

In previous work we also noted the relation between Q-scores and atomic B-factors (Pintilie & Chiu, 2021), and here we further explored and showed examples of how Q-scores can be converted to B-factors at resolutions between 1 and 4Å. We showed that when these B-factors are used to generate a model-map, the model-map is more similar to the experimentally-obtained 3DEM map than when not using atomic B-factors (or setting atomic B-factors to 0). Such estimated B-factors are useful annotations for 3DEM atomic coordinates archived in the PDB. We noted that the atomic B-factors discussed here are different from two other B-factors often mentioned in 3DEM: B-factors for sharpening a 3DEM map (Terwilliger *et al*., 2018), and the Rosenthal-Henderson B-factors to estimate the number of particles needed for certain resolution as constrained by instrumental and sample conditions (Rosenthal & Henderson, 2003).

To assess 3DEM entries in the EMDB (wwPDB Consortium, 2024), we also described here two percentile-based metrics: Q-relative-all and Q-relative-resolution. The Q-relative-all metric represents the overall quality of the map and model, comparing their Q-score to the entire EMDB archive. The higher the Q-relative-all-metric is, the higher the quality of the map and model. On the other hand, Q-relative-resolution compares the Q-score of a map and model to Q-scores of other entries of similar resolution. For this score, the closer it is to 50%, the more it is ‘as commonly observed’ for other entries in the EMDB of similar resolution. Q-relative-resolution scores that much higher (e.g. above 95%), or much lower (e.g. less than 5%) could indicate inconsistencies such as poorly fit models, oversharpened maps, overfitted models, or reported resolutions that may not fully reflect the entire map.

Finally, we note that Q-scores do not evaluate the stereochemical quality of an atomic coordinate model, such as proper bond lengths, bond angles, dihedral angles, chiral centers, etc. These attributes can be evaluated with other methods such as Molprobity (Williams *et al*., 2018). Within the EMDB and wwPDB OneDep system, the same methods are used to assess structures determined using 3DEM, MX, and nuclear magnetic resonance spectroscopy, to support deposition and rigorous validation(Gore *et al*., 2017; Feng *et al*., 2021; Young *et al*., 2017). We hope that Q-scores will continue to serve as a complementary and necessary metric alongside such other metrics, to reflect 3DEM map-model fit and map quality.

## Acknowledgements

The research was partially supported by the National Institutes of Health (R24GM154186).

Molecular graphics and analyses performed with UCSF Chimera and UCSF ChimeraX, developed by the Resource for Biocomputing, Visualization, and Informatics at the University of California, San Francisco, with support from NIH P41-GM103311, National Institutes of Health R01-GM129325 and the Office of Cyber Infrastructure and Computational Biology, National Institute of Allergy and Infectious Diseases.

RCSB PDB Core Operations are funded by the U.S. National Science Foundation (DBI-2321666, P.I.:S.K. Burley), the U.S. Department of Energy (DE-SC0019749, P.I.:S.K. Burley), and the National Cancer Institute, National Institute of Allergy and Infectious Diseases, and National Institute of General Medical Sciences of the National Institutes of Health under grant R01GM157729 (P.I.:S.K. Burley).

EMDB is supported by the European Molecular Biology Laboratory, European Bioinformatics Institute. EMDB is further supported by funding from the Wellcome Trust [212977/Z/18/Z]

## Data Availability

The Q-score method and code is available in the following GitHub repositories: https://github.com/gregdp/mapq https://github.com/gregdp/chimerax-qscore

Files with Q-scores and reported resolution for the plots generated here can be found in the same repository: https://github.com/gregdp/mapq/tree/master/data

## Appendix A. Methods

### A1. Q-score distributions

Q-scores were calculated using the Q-score plugin for UCSF Chimera v1.9.7. The only parameter that can be varied, sigma, which corresponds to the width of the reference Gaussian, was set to 0.4. Polynomial regression of Q-scores and reported resolution was performed in Microsoft Excel. Cross-correlation and Cross-correlation about the mean were performed using the function overlap_and_correlation in the FitMap module, in UCSF Chimera(Pettersen *et al*., 2004).

### A2. Residual Plots

R (https://www.r-project.org/) was used to further analyze the goodness of fit of the polynomial regression calculations and their appropriateness. Residual standard error was generated for each regression calculation, so were the residue plot and QQ-plot of the residuals. Examples of residual plots for average Q-scores of all maps and models, and maps and models with CC>=0.7 are illustrated in Figure S1. The residual plots and QQ plots were used to verify that the polynomial regression calculations are appropriate statistical models that both satisfy the assumptions and sufficiently describe the data fitting.

### A3. CC and CC-mean scores

Two formulations of the real-space cross-correlation scores were used: cross-correlation (CC) and cross-correlation about the mean (CC-mean).

The cross correlation (CC) score is defined as:

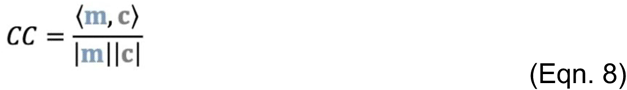

In Eqn. 8, <> denotes the inner product of two vectors. The vector **m** contains values taken from the grid points in the model-map. A model map is generated from atom coordinates using the Chimera *molmap* function, which takes two parameters: resolution, and grid spacing. Here, for resolution we used the reported resolution of the 3DEM map associated with the model, and for the grid spacing the default, resolution/3. The *molmap* function simply places a Gaussian at each atom position, with height proportional to the element number of the atom, and sigma proportional to the resolution, *d*: σ = d/(pi*sqrt(2)). The values in **m** are taken from the model map, but only at grid points where the values are above a threshold, here we used 0.01. The vector **c** contains values interpolated in the 3DEM map at spatial positions corresponding to the grid points of **m**.

Using the same vectors **m** and **c**, the cross-correlation about the mean (CC-mean) is calculated as:

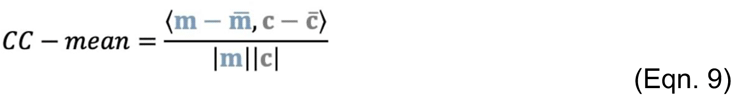

In Eqn 9, letters **m** and **c** refer to the same vectors as described above. When they have a bar above them, they are vectors with the same size but for which each entry is the average value of the respective vector.

### A4. Estimation of B-factors with Q-scores

B-factors were generated from Q-scores using Eqn. (5). A range of scaling factors, *f*, were applied, ranging from 0 to 300, in steps of 20. For each scaling factor, a model was generated with B-factors according to the scaling Eqn. (5), and then converted to a model-map using phenix.fmodel and phenix.mtz2map. The commands and parameters are as follows:

command 1: phenix.fmodel high_resolution=[map resolution] scattering_table=electron generate_fake_p1_symmetry=True [model file path]

command 2: phenix.mtz2map high_resolution=[map resolution] include_fmodel=true scattering_table=electron [model file path] [mtz file generated by command 1]

We set the [map resolution] parameter to be the same as the reported resolution of the 3DEM map to which the model-map is being compared. The resulting model-maps were then opened with Chimera. All points with values above 0.01 were considered, and the values at these points stored in a vector **m**. Values at corresponding positions were interpolated from the 3DEM map and stored in another vector **c**. The cross-correlation about the mean (CC-mean) between the vectors **m** and **c** was then calculated using the chimera.overlap_and_correlation function as per Eqn. 9.

### A5. Correlation between Q-relative-resolution and resolution

The correlation coefficient between Q-relative-resolution and the reported resolution is calculated as follows:

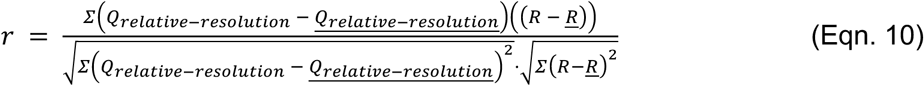

In Eqn 10, Q-relative-resolution is a vector with Q-relative-resolution values, while R is a vector with corresponding reported resolution values. Here, *<u>Q_relative-resolution_</u>* and <u>*R*</u> correspond to the mean values of Q-relative-resolution and resolution, respectively. The correlation coefficient r is calculated separately for two groups based on resolution: one group includes observations with a resolution smaller than 5 Å, and the other comprises those with a resolution of 5 Å or greater. This approach allows for the assessment of correlation across different resolution ranges.

### Supporting Information

**Supplementary Figure S1.**
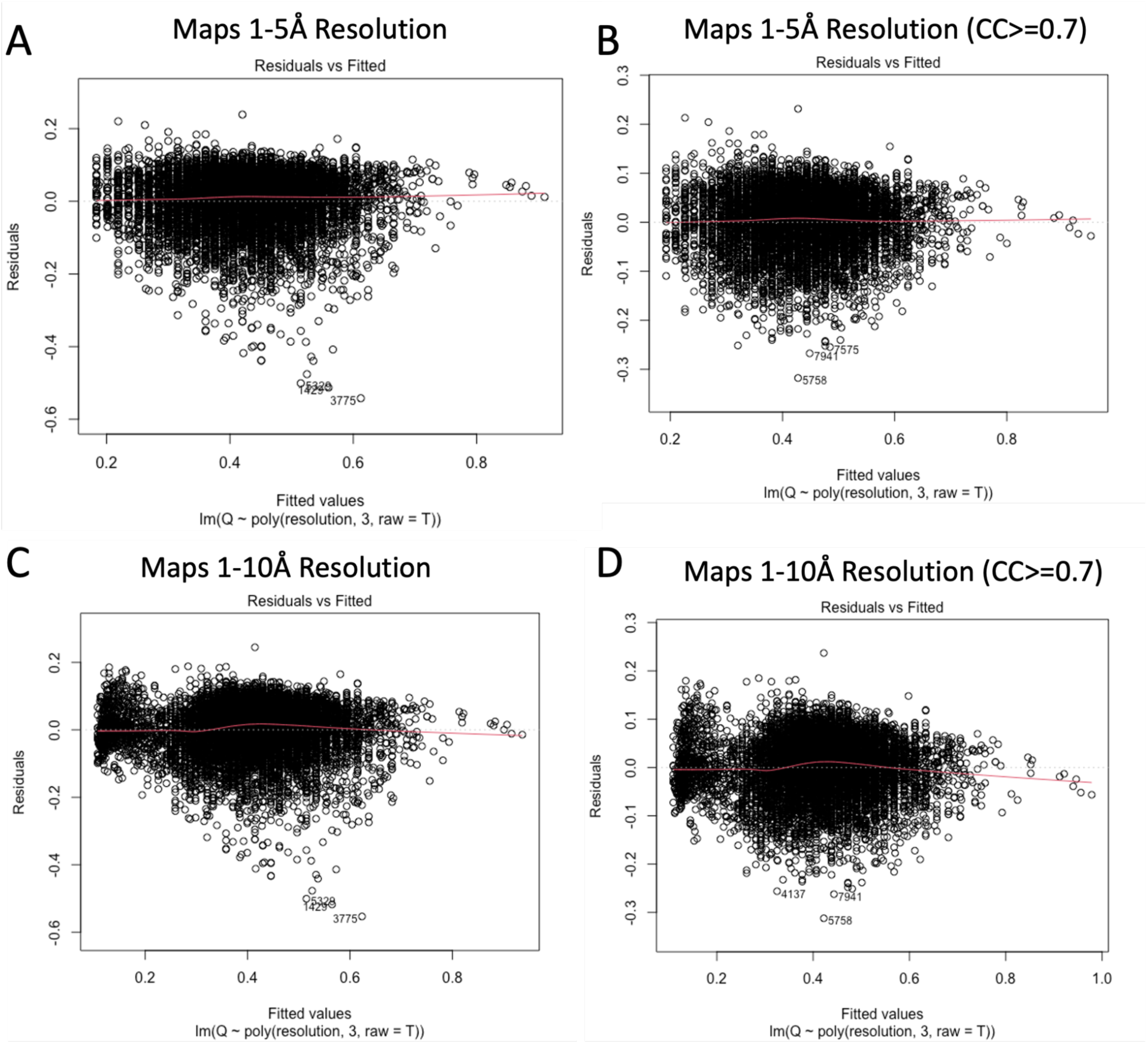
Residual plots for the regression calculation in Figure 1 of average Q-score vs. Reported Resolution, for (A) map-model pairs at 1-5Å resolution and (B) map-model pairs at 1-5Å resolution with map-model CC above 0.7. Residuals are well-distributed indicating the regression calculation is appropriate for this resolution range. Residual plots for the same regression calculations, but with maps in the resolution range 1-10Å are shown in (C) for all maps, and in (D) for maps with map-model pair CC above 0.7. An informative regression calculation should produce an unbiased and homoscedastic residual plot without any obvious pattern. Plot (B) indicates the best regression calculation among the four, and the other three plots demonstrates minor deviation from a perfect regression calculation, with plot (A) impacted by the outliers, and plots (C) and (D) impacted by the relatively different distribution for the resolution ranges of 1-5Å and 5-10Å. The residual standard errors are 0.075 for all maps with resolution 1-5 Å (A), 0.061 for maps with resolution 1-5Å and map-model pair CC >= 0.7 (B), 0.074 for all maps with resolution 1-10 Å (C), and 0.062 for maps with resolution 1-10Å and map-model pair CC>=0.7 (D). Both X and Y axes are scaled.

#### S1. Q-score distributions with outlier removal

While many map-model Q-scores appear close to the polynomial regression curve in Figure 1A, there are also some that are quite far from it. We tested whether removing some maps and models would change the correlation between Q-scores and reported resolution, aiming to remove maps and models that do not match well due to reasons such as inaccurate modeling. We applied two model-map metrics which measure the similarity between the 3DEM map and a model-computed map: cross-correlation about the mean (CC-mean) and cross-correlation (CC); these are further described in Methods. These two metrics are used because they can potentially identify which models are not fitted properly to the map(Pintilie & Chiu, 2012). Higher scores mean that the model matches the map better, and that it fits well. Lower scores mean the model does not match the map as well, and could potentially be incorrectly fitted.

Figures S1A-C show plots of Q-score vs. reported resolution before and after removing maps-model pairs with CC-mean<0.5 and CC<0.7. Different thresholds were used for CC-mean (threshold = 0.5) and CC (threshold = 0.7) in order to remove a similar number of entries from the datasets (∼1,000 entries). The chosen threshold values are different since CC scores tend to be higher than CC-mean scores. This is because when subtracting the mean for the CC-mean score, map values are lower, some becoming negative. The remaining map-model pairs have Q-scores much closer to the fitted polynomial regression curve, as plotted in Figures S1B and S1C. Figure S2E and S2F show Q-scores vs. resolution for the removed map-model pairs. When map-model pairs are removed because of CC-mean<0.5, the remaining map-model pairs have Q-scores mostly below the regression curve, while for the map-model pairs removed based on CC<0.7, many are close to the curve as well. The regression plots in Supplementary Figures S1H and S1I show that CC-mean scores are weakly correlated to Q-scores (R^2^=0.3264), while CC scores appear to have no correlation (R^2^=0.0071).

The plot for a small dataset with 386 map-model pairs from the EMDB/PDB reported previously(Burley *et al*., 2022) is shown in Supplementary Figure S2D. The regression curves for these 386 map-model pairs, for the data set of ∼10k map-model pairs, and for the data sets after removal of low CC and CC-mean scores are all plotted together for comparison in Supplementary Figure S2G, showing they are all very similar. Thus, the relationship between Q-score and reported resolution appears to be robust for different but representative data sets, and after removing some map-model pairs with low similarity (using CC-mean and CC scores).

**Supplementary Figure S2.**
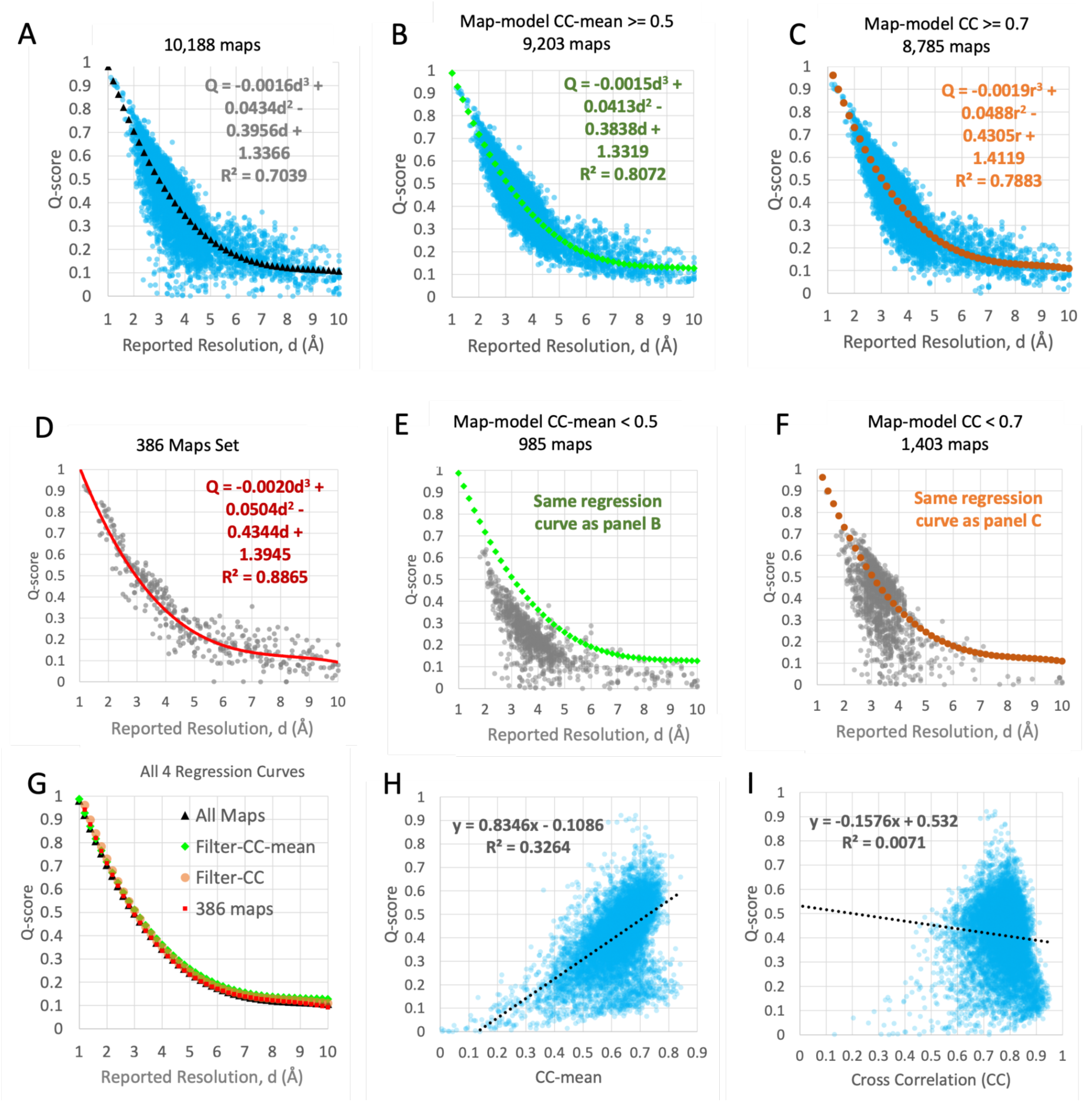
Regression analysis of Q-scores vs. reported resolution for 4 data sets (A-D). (A) 10,189 map-model pairs used in this analysis; note that this is the same dataset shown in Figure 1A. (B) Same data set as (A), but only maps and models with CC-mean >= 0.5. (C) Same as (A) but only maps and models with CC >= 0.7. (D) 386 maps-model pairs previously reported. (E) Map-model pairs with CC-mean < 0.5, with regression curve from panel B. (F) Map-model pairs with CC < 0.7 with regression curve from panel C. (G) All 4 regression curves for (A-C) and (D) plotted together. (H, I) Q-score vs. CC-mean and CC, with linear regression lines.

#### S2. Q-score Distribution Around Polynomial Regression Curve

We further characterize the distributions of Q-scores above and below the 3rd degree polynomial regression curve, with the aim of identifying outliers which are far from this regression line. We looked at how many maps and models have Q-scores within a small window of the curve when it is moved by an offset along the vertical Q-score axis. This offset acts to move the curve up and down. At each offset, we count the number of map-model pair data points close to the curve at the offset position, within a small window size, e.g. w=0.01, as illustrated in Supplementary Figure S3A. Supplementary Figure S3B plots the number of maps and models for offsets in the range [−0.3, 0.3]. This range of offset was used because beyond +/−0.3, there are very few Q-scores around the offset curve. The plot in Supplementary Figure S3B shows that the 0-offset regression curve, which represents the mean or average Q-score, is slightly below the median which is found at offset=0.012, and the peak which is found at offset=0.024. The median represents the offset at which half of the maps have Q-scores below the curve and the other half have higher Q-scores, while the peak represents the offset at which the highest number of maps have corresponding Q-score.

**Supplementary Figure S3.**
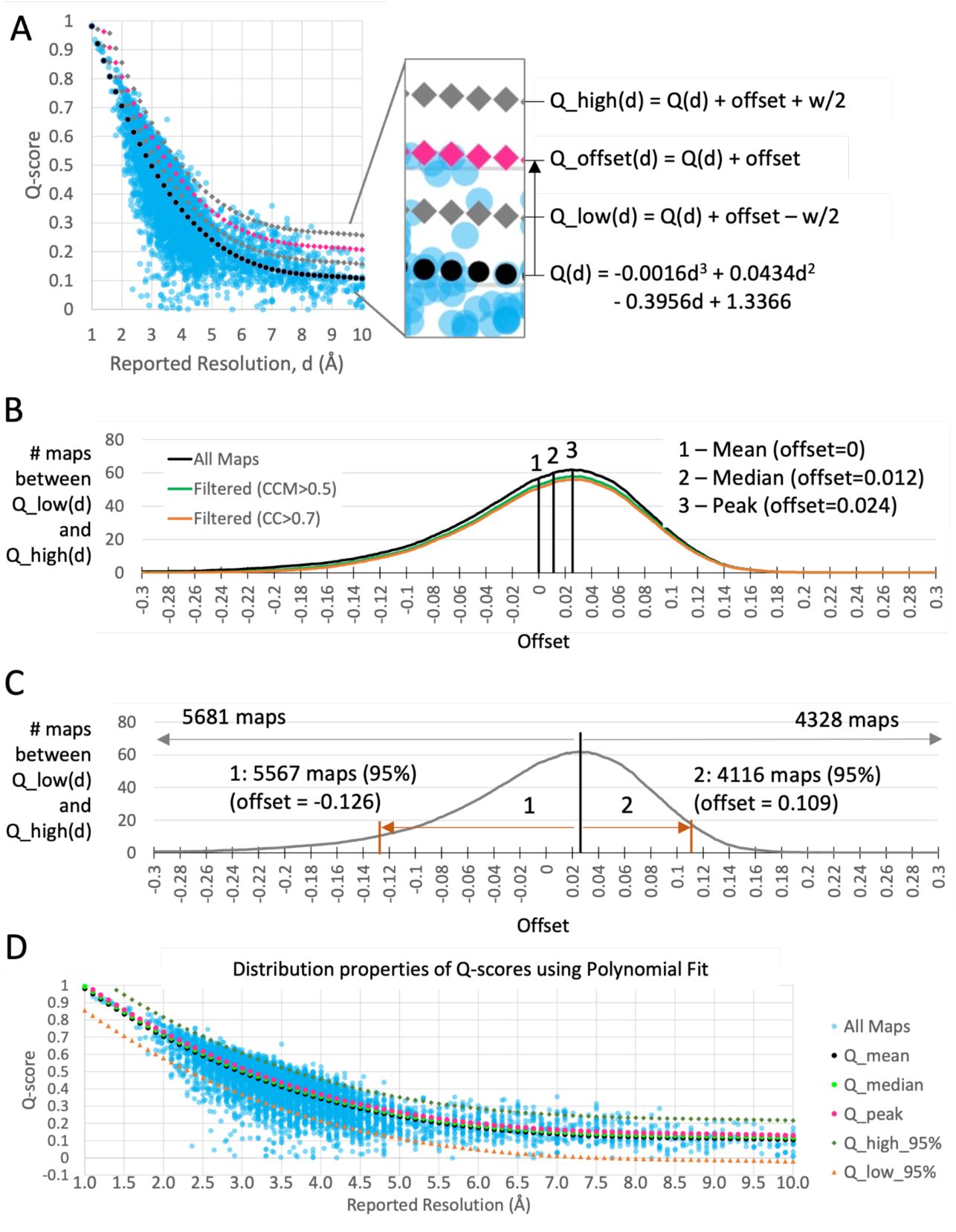
Distribution of Q-scores based on the 3rd degree polynomial regression curve. (A) The regression curve for Q-scores vs. reported resolution, Q(d), is plotted with a dotted black line. The same curve is also plotted with an offset, and upper and lower bounds using window size w. (B) The number of maps with Q-scores within a window of size w around the offset is plotted (w=0.01), vs. the offset ranging between [−0.3,0.3]. Q-scores of the mean, median, and peak are marked on the plot. (C) Two offsets are marked on the curve which enclose 95% of the maps around the peak Q-score value. Note that plots in B and C include all data points. (D) The regression curves representing mean, median, and peak are plotted. Curves enclosing top and bottom 95% of the maps above and below the peak are also plotted.

Supplementary Figure S3B shows that the Q-score distribution around the polynomial regression curve is close to a normal distribution, although it is left-skewed, i.e. it decreases more slowly on the left side. The top and bottom offsets on either side of the peak which enclose 95% of the data are marked in Supplementary Figure S3C. The regression curve is plotted with corresponding offsets for the mean, median, and peak in Supplementary Figure S3D, which are very similar, and at the offset values enclosing 95% of the data.

#### S3. Q-score Distribution Using Rolling Window

We further investigated Q-score distribution using a rolling window over the resolution range 1.0 to 10 Å. This allows us to examine Q-score distribution at each resolution without the need of a regression curve fitted to all the data points. At each resolution *d*, maps and models with resolution *d*-w/2 to *d*+w/2 are considered, as illustrated for resolution *d*=2.5 Å in Supplementary Figure S4A. The number of maps and models with resolution within this range vs. Q-score is plotted in Supplementary Figure S4B. The curve is close to a normal distribution but also decreasing more slowly on the left (towards lower Q-scores). The mean, median, and peak are calculated for this distribution and shown on the plot in Supplementary Figure S4B. The top and bottom Q-scores within which 95% of the maps and models in this resolution range are included are also calculated and marked on the curve in Supplementary Figure S4B. This procedure was repeated for resolutions in the range 1-10 Å; all the peak and 95% thresholds at each resolution are plotted in Supplementary Figure S4C. They are similar to the smoother polynomial regression curves with corresponding offsets, as shown in Supplementary Figure S4D. The values obtained with the rolling window are however discontinuous, especially in the resolution range 1.0 to 2.0 Å, where there are fewer maps and models.

**Supplementary Figure S4.**
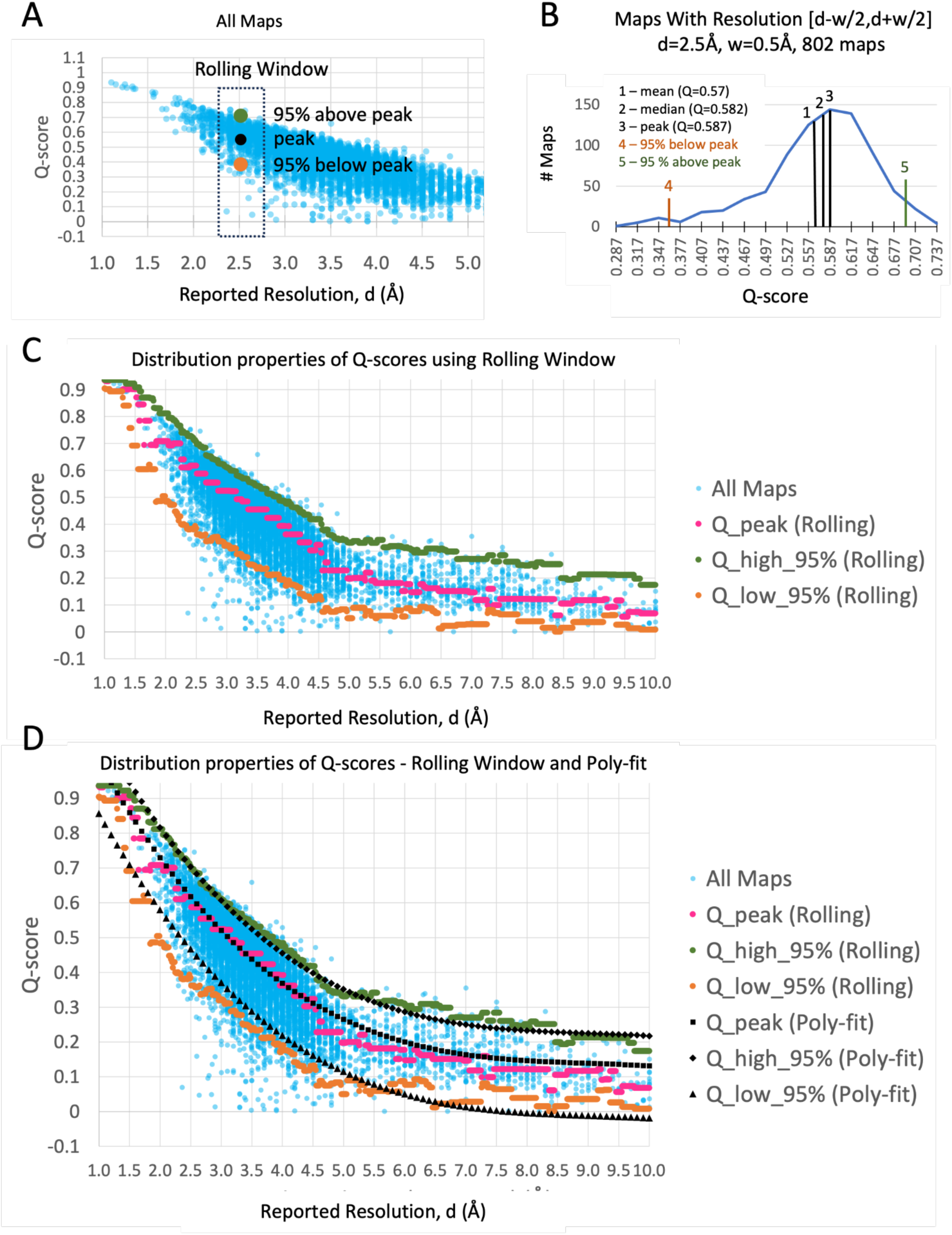
Calculation of mean, median, peak, and percentiles using rolling window approach. (A) Plot of Q-score vs. resolution for a subset of the ∼10k maps and models with resolution < 5.0Å. A window positioned at reported resolution d=2.5Å is shown, with peak and 95% percentile points. (B) Distribution curve for the 802 maps within a window of size w=0.5Å, with mean, median, peak, and 95% percentiles marked. (C) Plot of all ∼10k maps and models vs. resolution, showing the mean, median, peak, and 95% percentile points obtained at each resolution using rolling windows. (D) Comparison of rolling window (Rolling) results with the polynomial regression curves (Poly-fit).

**Supplementary Figure S5.**
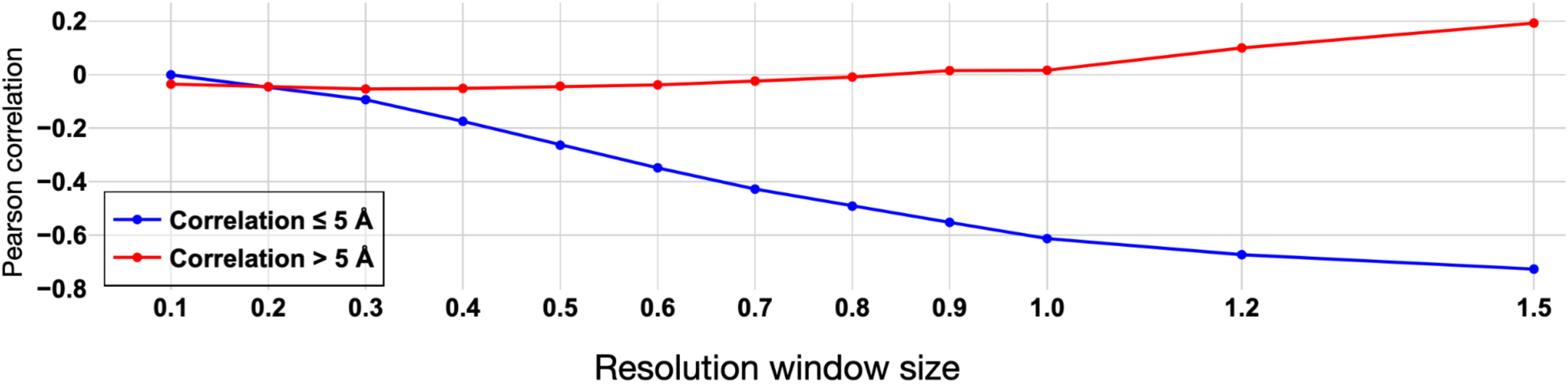
Plot of Pearson correlation between Q-relative-resolution and reported map resolution vs. resolution window sizes. The blue curve is for maps in EMDB with reported resolution ≤ 5 Å, while the red curve is for maps in EMB with reported resolution > 5 Å.

## Notes

https://github.com/gregdp/mapq

https://github.com/gregdp/chimerax-qscore

### Competing Interest Statement

The authors have declared no competing interest.

